# Refresh my memory: Episodic memory reinstatements intrude on working memory maintenance

**DOI:** 10.1101/170720

**Authors:** Abigail N. Hoskin, Aaron M. Bornstein, Kenneth A. Norman, Jonathan D. Cohen

## Abstract

A fundamental question in memory research is how different forms of memory interact. Previous research has shown that people rely on working memory (WM) in short-term recognition tasks; a common view is that episodic memory (EM) only influences performance on these tasks when WM maintenance is disrupted. However, retrieval of memories from EM has been widely observed during brief periods of quiescence, raising the possibility that EM retrievals during maintenance-critically, before a response can be prepared-might affect short-term recognition memory performance even in the absence of distraction. We hypothesized that this influence would be mediated by the lingering presence of reactivated EM content in WM. We obtained support for this hypothesis in three experiments, showing that delay-period EM reactivation introduces incidentally-associated information (*context*) into WM, and that these retrieved associations negatively impact subsequent recognition, leading to substitution errors (Experiment 1) and slowing of accurate responses (Experiment 2). fMRI pattern analysis showed that slowing is mediated by the content of EM reinstatement (Experiment 3). These results expose a previously hidden influence of EM on WM, raising new questions about the adaptive nature of their interaction.

## Introduction

Our memories do not exist in isolation, and neither do the neural circuits that represent them. Experiences may produce transient records in working memory — a temporary store for information to be maintained and manipulated over delays of seconds (Baddeley 1992; Baddeley & Hitch 1974; Repov & Baddeley 2006). Experiences can also simultaneously lay down more lasting traces as episodic memories, available to be recalled at a later time (beyond minutes), allowing us to relive specific, previously experienced events tied to the time and place of their occurrence (Tulving, 1983).

Early models proposed that working memory and long-term memory operated wholly in parallel (Shallice & Warrington, 1970). Evidence for the dissociation between working memory and episodic memory largely came from lesion studies, which found that damage to the medial temporal lobe (MTL) caused severe episodic memory deficits (Cave & Squire, 1992; Squire, 1992), while working memory, associated with the prefrontal cortex (Cohen et al., 1994), remained intact (Drachman & Arbit, 1966). More recent models propose that they support each other (Cohen & O’Reilly, 1996; Baddeley, 2000). There is accumulating evidence that episodic memory, and its neural substrates in the MTL, are engaged during short-term memory tasks that also engage working memory (Lewis-Peacock et al., 2016; Ranganath, 2005; Ranganath et al. 2004, 2005; Ranganath & Blumenfeld 2005; Axmacher et al. 2007), suggesting these memory systems do not operate entirely independently of one another.

Experiments testing for an interaction between these two types of memory have largely focused on the hypothesis that episodic memory is used to support working memory when maintenance is disrupted, leading to errors that reflect features of episodic memory. For instance, participants show proactive interference from recently studied stimuli when working memory is disrupted for 18 seconds (Wickens et al., 1976). Here, we ask whether episodic memory *only* contributes when working memory is disrupted, or whether it contributes more ubiquitously.

## Do ongoing reinstatements from episodic memory influence working memory, even in the absence of distraction?

A growing number of studies indicate that, during periods of rest, the neural structures that support episodic memory (EM) are active (Buckner, 2010) and appear to be reinstating recent experiences (Tambini et al., 2010) or activating potential future scenarios constructed on the basis of past experiences (Buckner & Carroll, 2007). These reinstatements trigger coordinated activity patterns across a broad swath of cortical regions, including those presumably involved in working memory (WM) maintenance, such as prefrontal cortex (Miller & Cohen, 2001). This widespread activation is reliably present even during brief lapses in external stimulation (Logothetis et al., 2012), such as those typically used as maintenance periods in WM experiments.

These observations lead us to ask the question: Do ongoing reinstatements from episodic memory affect the content of working memory, even when the latter is not being disrupted? Such an effect has not previously been observed — indeed, the lack of such an observation has encouraged researchers to employ these distraction-free maintenance tasks as specific probes of WM alone. However, previous studies measured only accuracy (e.g., proportion of substitution errors). We hypothesized that an influence of EM on WM search might be observable by other, more sensitive means: through examination of reaction times, and through the use of content-specific pattern analysis in neuroimaging.

## Using context as a signature of EM

To test our hypothesis, we leverage the fact that retrievals from EM carry with them temporal and associative context (Howard & Kahana, 2002), such that triggering the recall of one memory from a given context can cause the subsequent, involuntary recall of other memories sharing that context (Hupbach et al. 2009; Bornstein & Norman, 2017). This can occur even at the short delays typically associated with WM (Hannula et al., 2006). Therefore, we reasoned that — if reinstatements from EM occurred during WM maintenance — these reinstatements would likely be of memories that shared an encoding context with the target stimuli. Even if these reinstated memories do not lead to overt errors, they may intrude on or degrade other, task-relevant representations being maintained in WM, and thereby affect search and response times on subsequent decisions — even several seconds later, and even in the absence of further EM reinstatement. They may also express themselves in patterns of neural activity reflective of the reinstated memories.

## Present study: Three experiments measuring how EM reinstatements can inject contextual associates into WM, even in the absence of distraction

We present three experiments testing the hypothesis that context reinstated from EM intrudes on WM maintenance. In Experiment 1, we show that participants substitute same-context items in response to interference in a classic short-term delayed recall task with distraction during the maintenance period. These intrusions are distinct from the recency effect traditionally used to identify episodic influence in this task. In Experiment 2, we show that the influence of reinstated context is evident in response times, even when accuracy is at ceiling. In Experiment 3, we repeat the task from Experiment 2 with fMRI, and use multivariate pattern analysis (MVPA) to generate a trial-by-trial neural measure of how likely it was that participants were recalling a specific past context. We use this neural index of reinstatement to predict the degree of response time bias on a given trial. Finally, we show that EM reinstatement affects responses via a specific effect on the contents of WM.

Together, the results of these experiments reveal a novel aspect of the interaction between EM and WM: When target items are stored in WM, ongoing reinstatements from EM can inject contextual associates of these targets into WM, leading to confusion about whether these associates were part of the target set.

## Experiment 1

Previous studies using short-term recall tests have found that distraction during delay periods causes participants to rely on episodic rather than WM, as evidenced by the fact that errors are primarily words substituted from recent trials (Brown, 1958; Peterson & Peterson, 1959; Rose et al., 2014; Lewis-Peacock et al., 2016; Zanto et al., 2016). Here, we tested whether these substitutions can be biased by the encoding context of the target words. Specifically, if the four target words are sampled from one of the 12-word encoding contexts established at the outset of the experiment, does this lead to substitution of other (non-target) words from the same context? The logic of the study is shown in Figure 1, and examples of the initial context learning and delayed recall trials are shown in Figures 2A and 2B, respectively.

**Figure 1.**
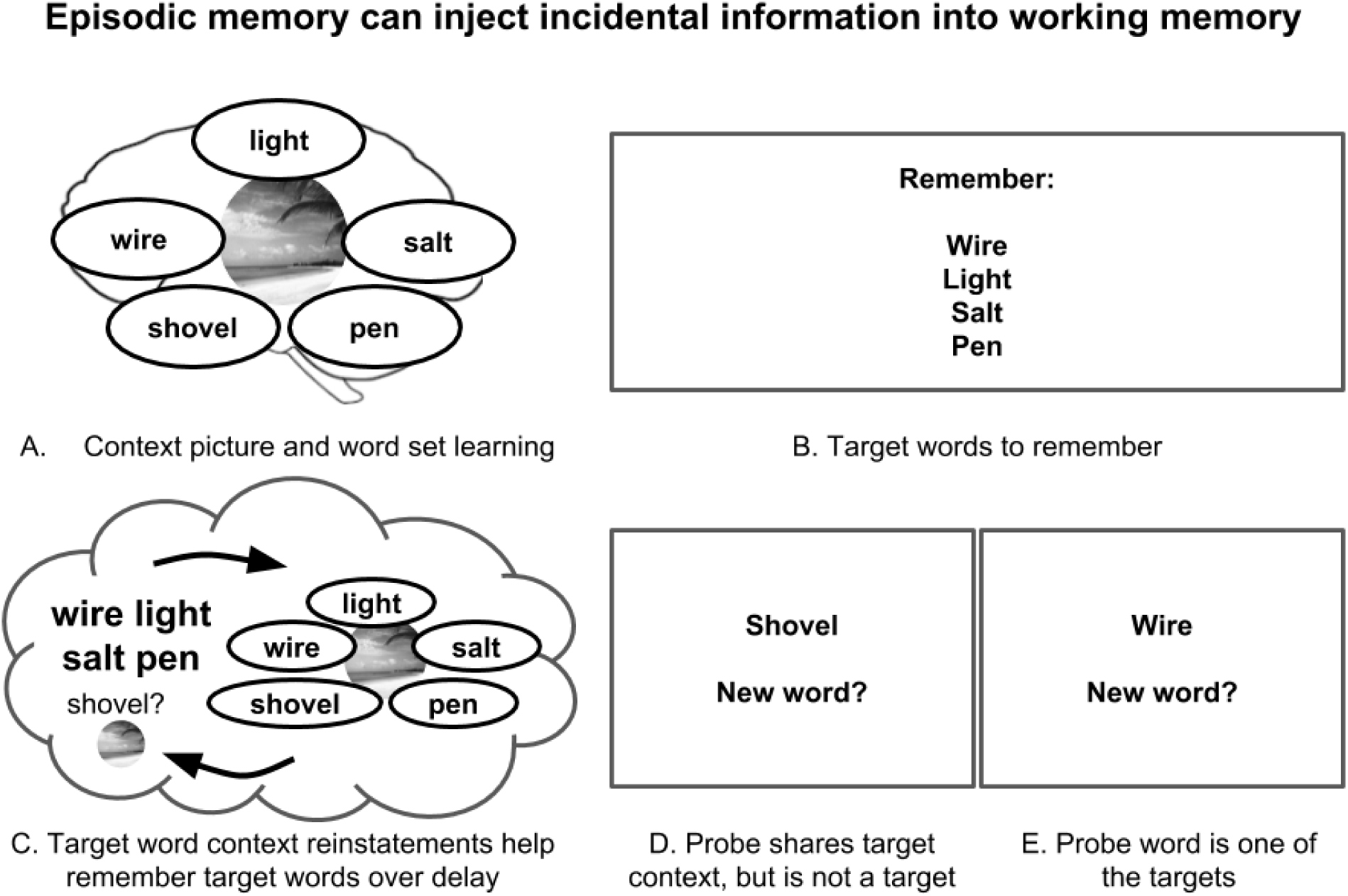
Episodic memory can inject incidental information into working memory. **A.** Episodic memory encodes items along with the context in which they were learned. **B.** When presented with target items to maintain over a delay period, working memory maintenance may be periodically influenced by reinstatements from episodic memory. **C.** These reinstatements may contain other items sharing the encoding context of the target items. **D.** These items might affect subsequent behavior, by impeding decision making when these items support the incorrect decision; **E.** and / or by facilitating decision making when they support the correct decision.

**Figure 2.**
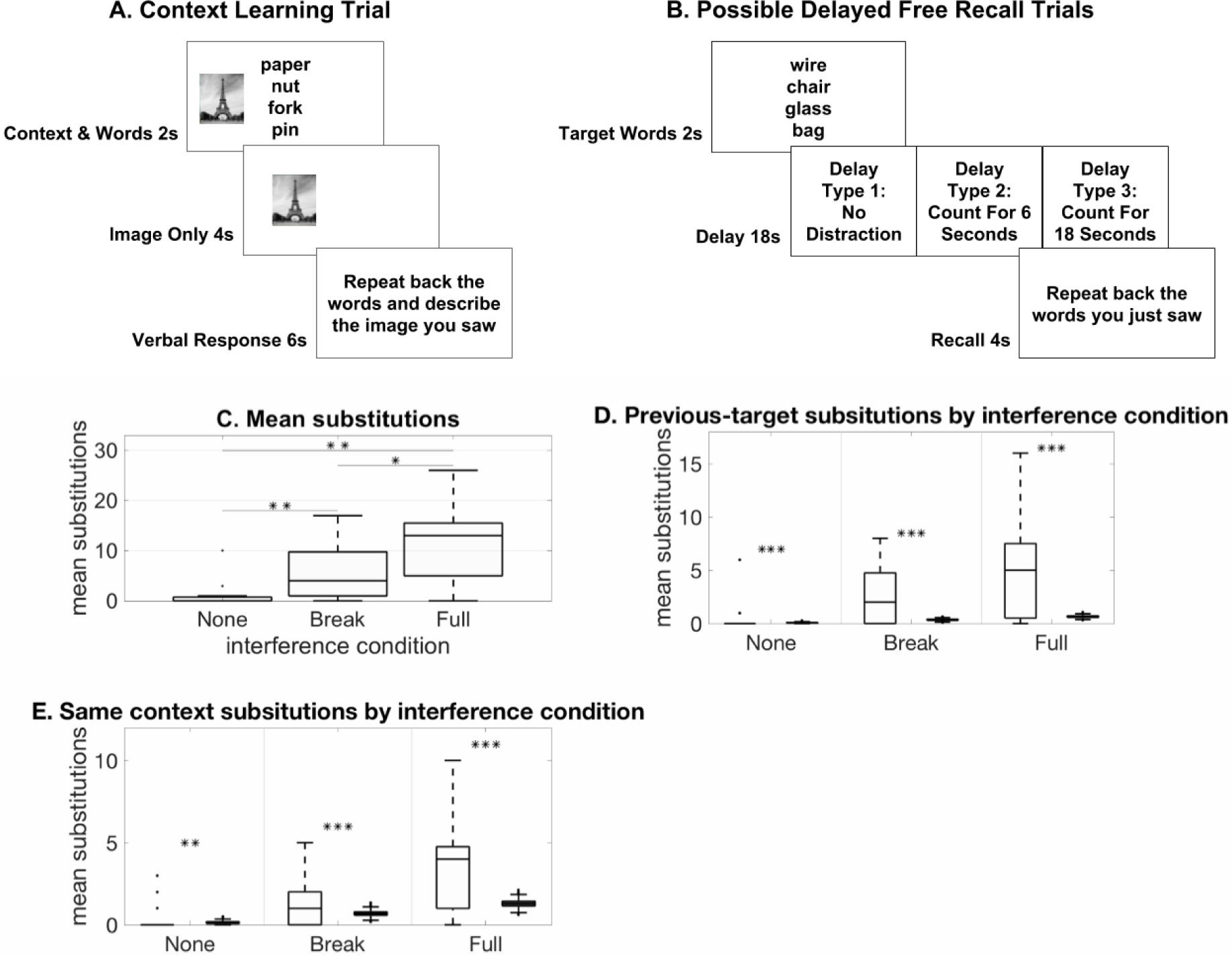
Experiment 1: Free recall task with added context. **A.** Participants (n = 15) studied lists of words in contexts distinguished by different pictures. **B.** We probed how these contexts affect performance on a short term recall task under three conditions: 1. when working memory was not disrupted, 2. briefly disrupted (break distraction), or 3. completely disrupted (full distraction). **C.** Participants made more errors in the full distraction than the break distraction condition (*t*(14) = 3.2756; *p* < .01; paired, two-sided t-test), and more errors in the full distraction than the no distraction condition (*t*(14) = 6.4526, *p* < .001; *p* < .01; paired, two-sided t-test). Participants also made more errors in the break distraction condition than the no distraction condition (*t*(14) = 4.4852, *p* < .001; *p* < .01; paired, two-sided t-test). * signifies *p* < .05, ** signifies *p* < .01, *** signifies *p* < .001. Black horizontal lines within boxes indicate median substitutions. Bottom and top edges of the box indicate the 25th and 75th percentiles. Whiskers extend to the most extreme data points not considered outliers. Black points outside boxes indicate outliers. **D.** Within each interference condition, left bars reflect subject data and right bars reflect simulated data based on randomized substitutions from the experiment’s word set. In all three conditions, participants made errors that reflected the influence of reinstated context. Specifically, participants substituted words from the previous trial at a higher rate than would be expected if they were randomly substituting words previously learned in the experiment. As computed by bootstrap analysis, the number of previous trial substitutions was greater than chance on full interference (*p* < .001), break interference (*p* < .001), and no interference trials (*p* < .001). This suggests that information from previous trials from episodic memory entered working memory, even when working memory was not overloaded. **E.** Participants also made substitution errors during recall that reflected the encoding context of the target set, or *same context* errors, at a higher rate than would be expected if they were randomly substituting words previously learned in the experiment. As computed by bootstrap analysis, the amount of same context errors made was greater than chance on full interference (*p* = .001), break interference (*p* = .001), and no interference trials (*p* = .025). This suggests that context information from episodic memory entered working memory, even when working memory was not overloaded. Box plots follow the same conventions as in D.

### Methods and materials

#### Participants

15 Princeton psychology students (9 females; ages 18 to 22) completed the study for course credit. All participants had normal or corrected-to-normal vision and provided informed consent. The study protocol was approved by the Princeton University IRB.

#### Stimuli

The experiment used six scene pictures, each of which served as a “context” that uniquely linked one of six sets of 12 words. The pictures were color photographs of outdoor scenes. The words were concrete nouns drawn from the Medical Research Council Psycholinguistic Database (Wilson, 1988). All words had a maximum of two syllables, Kucera-Francis written frequency of at least 2, a familiarity rating of at least 200, a concreteness rating of at least 500, and an imageability rating of at least 500. Pictures consisted of famous landmarks without any people in them. The words used in each set and the image associated with each set were randomized across participants.

#### Procedure

##### Word-context learning trials

The goal of the initial *context learning* phase was to associate words with distinct encoding contexts. On each of 48 learning trials, participants were shown four words drawn from the same set alongside the photograph associated with that set (Figure 2A). The picture served as an *encoding context*. To help participants encode the 12 words associated with the same picture as all belonging to the same context, each word was presented three times along with three other words randomly sampled from the same set and always displayed in the same context (i.e., with the picture associated with that list). On each trial, the four words and the picture associated with those words were presented for two seconds before the words disappeared and the picture remained on-screen. Four seconds later, the context picture was replaced by a prompt asking participants to vocally repeat back the four words just shown, and to then briefly describe the picture they had just seen. Participants were given six seconds to respond. Trials were of fixed length, regardless of participant’s responses.

##### Free recall phase

After the learning block was completed, participants performed 54 trials of a short-term retention task. On each trial, participants were shown four *target* words. The four target words were all drawn from the same context. No picture was presented alongside the words. Words remained on the screen for two seconds and were followed by an 18 second delay.

There were three types of delay (Figure 2B). Delay trial types were randomly intermixed, with 18 trials of each type. In the *no distraction* condition, participants were shown a fixation cross, in the center of the screen, for the entirety of the 18 second delay. In the *break distraction* condition, participants were shown a fixation cross in the center of the screen for six seconds. After six seconds, participants were shown a randomly generated three-digit number in the center of the screen. The number served as a prompt to count down out loud by sevens, starting at that number. After six seconds of counting, participants were again shown a centered fixation cross for six more seconds. In the *full distraction* condition, participants were shown a three-digit number at the start of the delay period, and instructed to count backwards out loud by sevens, starting from the prompted number, for the entire delay period.

In all conditions, participants were given eight seconds after the delay period to vocally recall the words shown at the beginning of the trial. These responses were recorded and scored for the number of words correctly recalled (zero through four). Mistakes were categorized as one of three types: 1. words from the same encoding context as the targets, 2. words from the previous free recall trial, or 3. other words learned during the experiment but not in categories 1 or 2. (No substitutions were made using words not learned during the experiment.)

### Experiment 1 results

We expected to see increasing numbers of substitution errors as the demands on working memory increased; therefore, we predicted participants would make the fewest substitutions following delays with no distraction, and the most substitutions following full distraction.

Consistent with our predictions, participants made more errors in the full distraction condition than in the break distraction condition (*t*(14) = 3.2756; *p* < .01; paired, two-sided t-test) and the no distraction condition (*t*(14) = 6.4526, *p* < .001; Figure 2C), and more errors in the break distraction condition than the no distraction condition (*t*(14) = 4.4852, *p* < .001; Figure 2C).

We also predicted that distraction would increase reliance on episodic memory and, accordingly, that substitution errors would reflect information retrieved from episodic memory. To test this hypothesis, we marked errors as belonging to one of three categories, two that specifically reflected intrusions from episodic memory: *previous-target substitutions* and *same-context substitutions;* as well as *other errors,* that reflected intrusions or failures of other kinds. These categories were motivated by the following considerations. First we expected recently-experienced words — in particular, the four words from the trial immediately previous — to be most accessible in episodic memory, and therefore likely to be recalled, brought into working memory, and mistakenly invoke a target response. We refer to these as *previous-target* substitutions. Second, we expected that maintaining target words in working memory would trigger episodic memory reinstatement of the context in which these words were studied (Howard & Kahana, 2002; Gershman et al., 2013). If this occurs, we should see an elevated substitution rate for the eight words that were studied in the same context as the target words, but that were not part of the current trial’s target set. We refer to these as *same context* substitutions. The context from which the target words were drawn changed with each trial, ensuring that previous-target and same context substitutions were mutually exclusive possibilities. Finally, we refer to substitutions from one of the 56 remaining words learned in the experiment, that were neither targets, *previous-target* or *same context* errors, as *other* errors.

By categorizing errors in this way, we could compare the number of each kind of error to the number that would be expected if the errors were drawn at random from the 68 possible non-target words. While all three kinds of words should be present in episodic memory, we predicted that *previous-target* errors, reflecting recency, and *same context* errors, reflecting the bias towards clustered recall of items sharing encoding context, should be overrepresented relative to *other* errors.

If substitution errors were uniformly distributed among the 68 possible words, only 4/68 of the errors made in each interference condition should be *previous-target* substitutions. Participants substituted words from the previous trial at a higher rate than would be expected if they were randomly substituting words previously learned in the experiment (Figure 2D). As computed by bootstrap analysis, the amount of previous trial substitutions made was greater than chance on full interference (subject mean = 5.20, std = 4.95; bootstrapped mean = .64, std = .10; *p* < .001), break interference (subject mean = 2.67, std = 3.04; bootstrapped mean = .34, std = .08; *p* <.001), and no interference trials (subject mean = .47, std = 1.55; bootstrapped mean = .07, std = .04; *p* < .001). This suggests that information from previous trials from episodic memory entered working memory, even when working memory was not overloaded.

Similarly, if substitution errors were uniformly distributed among the 68 possible words, only 8/68 of the errors made in each interference condition should be *same context* substitutions. Instead, on full interference trials, the proportion of *same context* substitutions was greater than what would be expected by chance (subject mean = 3.40, std = 2.77; bootstrapped mean 1.29, std = .21; *p* = .001). This suggests that context information was indeed affecting decision making when working memory was overloaded (Figure 2E). Same context substitutions were also greater than what would be expected by chance in the break condition (subject mean = 1.33, std = 1.59; bootstrapped mean = .86, std = .19; *p* = .001). Critically, although the frequency of substitutions on the no interference trials was low (mean = 1.13, std = 2.67; Figure 2C), when they did occur, they were biased towards coming from the same context as the target words (subject mean = .40, std = .91; bootstrapped mean = .13, std = .08; *p* = .025).

### Experiment 1 discussion

Participants completed a short term retention task with three distraction conditions. When there was no distraction during the retention delay, participants made almost no errors, consistent with the idea that they were able to easily use working memory to complete this task. Errors increased when participants were made to perform a distractor task midway through the delay, and were further increased when the distractor task spanned the entire retention interval. These errors took the form of substituting other words from the experiment in place of the current trial’s target words.

A disproportionate number of substitutions were made using words from the same encoding context as the target words, despite the fact that these kinds of words represented only a small fraction of the words used on the task. This distribution of substitutions is consistent with previous observations that, when working memory maintenance is interrupted, participants rely on recency-biased retrievals from episodic memory (Lewis-Peacock et al., 2016; Zanto et al., 2016; Rose et al., 2014). Critically, our results also establish that the context-based nature of errors can serve as an additional signature of episodic memory recruitment in these tasks, augmenting the suite of tools available to identify EM recruitment. As would normally be predicted, both kinds of errors were most evident when retention in working memory was subject to interference. Notably, however, the pattern of errors indicated the engagement of episodic memory even when distraction was momentary, hinting that it might be present even in the absence of distraction — that is, under conditions ordinarily assumed to rely exclusively on working memory.

Our findings raise two questions. First, does episodic memory affect working memory in the absence of external distraction? While substitutions in the no distraction condition were significantly biased toward being from the same encoding context as the target words, there were very few errors (of any kind) in this condition, making us wary of drawing strong conclusions from this result on its own. Second, when during the task does episodic memory retrieval occur, and how does it influence performance? Are episodic memories retrieved during the delay, either incidentally and/or to support maintenance, or strictly at the time of response? We use the signature of context effects established in Experiment 1 to address these questions in Experiments 2 and 3.

## Experiment 2

In Experiment 2, we used a more sensitive measure, reaction time (RT), to investigate the effect of context on behavior. Participants performed the same context training exercise from Experiment 1 (Figure 3A), this time followed by a delayed non-match to sample task (DNMS; Figure 3B) with no distractions during the delay periods.

**Figure 3.**
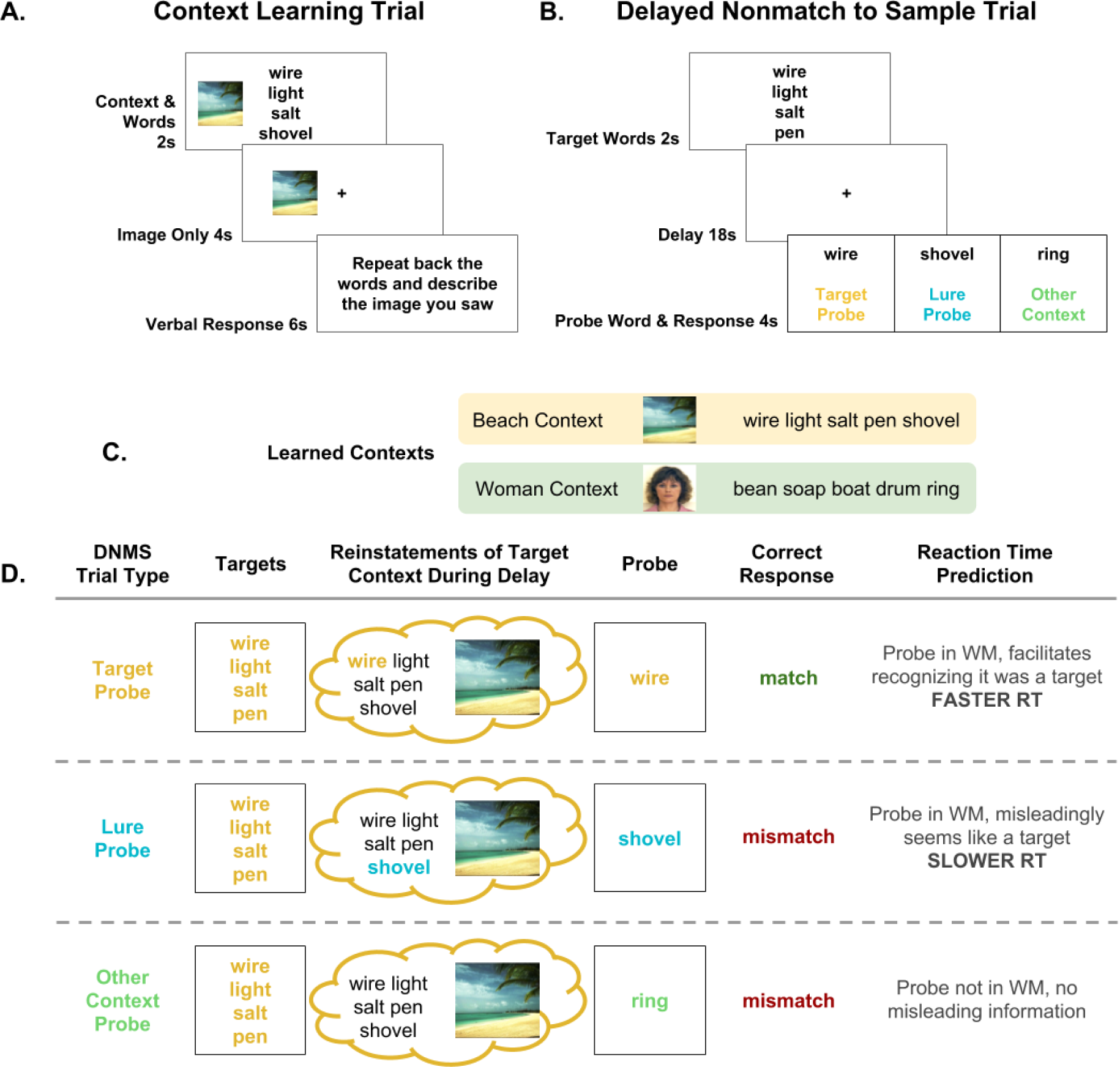
Experiment 2: DNMS task with added context. **A.** In the *context learning* phase, participants studied 48 words that were split into four sets of 12. Each set was paired with a unique *context picture*. **B.** In the *testing* phase, participants performed a delayed non-match to sample (DNMS) task, in which they remembered four *target* words across an 18 second delay. After the delay, they were shown a single *probe* word and asked whether that word was *not* one of the four they had just seen. Response times were recorded and used as a measure of whether the participants’ performance had been affected by context information reinstated from episodic memory. **C.** Subsets of two example contexts are presented for illustrative purposes. **D.** We hypothesized that the contents of working memory are influenced by reinstatements from episodic memory. These reinstatements activate working memory representations of trial-irrelevant words that were linked to the target words during the context learning phase. We predicted that, when the probe word was one of the targets, participants would be fastest to respond since the target probe should clearly match the content of working memory, allowing the search process to terminate quickly. For *non-target probe* trials, we predicted participants would respond more slowly since they needed to exhaustively search through the contents of working memory to decide to reject the probe. Within non-target probe trials, we predicted participants would be slowest to respond to *lure probes*, since these probes would match the context information in working memory elicited by the target words but mismatch the actual target words. Since this conflicting evidence was not present in *other context probe* trials — the probe word did not match the context information or target words in working memory — we predicted participants would be less impaired on *other probe* trials.

### Methods and materials

#### Participants

88 Princeton students (55 females; ages 18 to 21; native English speakers) completed the study for course credit. All participants had normal or corrected-to-normal vision and provided informed consent. The Princeton University IRB approved the study protocol. 8 participants were excluded from RT analyses on the basis of their accuracy scores being less than chance performance, leaving the participants reported here.

#### Procedure

In the *Learning phase*, participants studied four different sets of words, each containing 12 words drawn from the same set of words used in Experiment 1. Each word set was paired with a unique context picture. The paired words and orientation of each context picture were randomly assigned anew for each participant. *Learning phase* trials followed the same procedure as in Experiment 1 (Figure 2A; Figure 3A), now over four contexts of 12 words each.

In the *Testing phase*, participants performed 60 trials of a DNMS task, in which targets were selected from the words learned in the learning phase (Figure 3B). On each trial, one context was selected at random, and then four target words were selected from within that context. These words were shown on the screen together for two seconds — critically, without the associated context image. When the words disappeared, they were replaced by a centered fixation cross, displayed for 18 seconds. Participants were instructed to use this delay to remember the four words they had just seen. There was no distraction during the delay period.

After the delay period, participants were shown a probe word and asked to respond *mismatch* if the given word was not one of the four they had just seen on this trial, or match if it was one of the four target words. The keys used to signify *mismatch* and *match* — the left and right arrows — were counterbalanced across participants. A successful response was indicated by a green fixation cross while an unsuccessful response (incorrect response or time-out after four seconds) was indicated with a red fixation cross.

Probe words could be one of three types: 1. *target* probes were drawn from the four-word target set presented on the current trial; 2. *lure probes* were drawn from the same context list as the target words, but, critically, these probes were not one of the target words; 3. *other context* words were drawn from one of the three contexts other than the one from which the target words were drawn. Target probes were drawn from the target words, so the correct response to target probes was that they were a “match” to the targets; lure and other context probe words did not contain one of the target words, so the correct response on lure and other context probe trials was “mismatch”. Participants were not signaled as to which kind of probe was being used on each trial.

There were equal numbers of target, lure, and other context probe trials, so a participant who responded “mismatch” on every trial would be correct on 66% of trials. 8 participants fell below this accuracy threshold, whom we excluded from further analysis.

### Experiment 2 results

#### Accuracy

Given the absence of distraction, accuracy was high across all three conditions (mean = 94.84%, SEM = .78%) with no significant differences in accuracy between *target* (mean = 95.01%, SEM = .82%, *other context* (mean = 95.10%, SEM = .74%), or *lure* trials (mean = 94.31%, SEM = .78%), (*p* > .2 by paired, two-sided t-tests for all pairwise comparisons; Figure 4A). Because these inaccurate trials were rare and did not vary in proportion between categories, we excluded inaccurate trials from the RT analyses.

**Figure 4.**
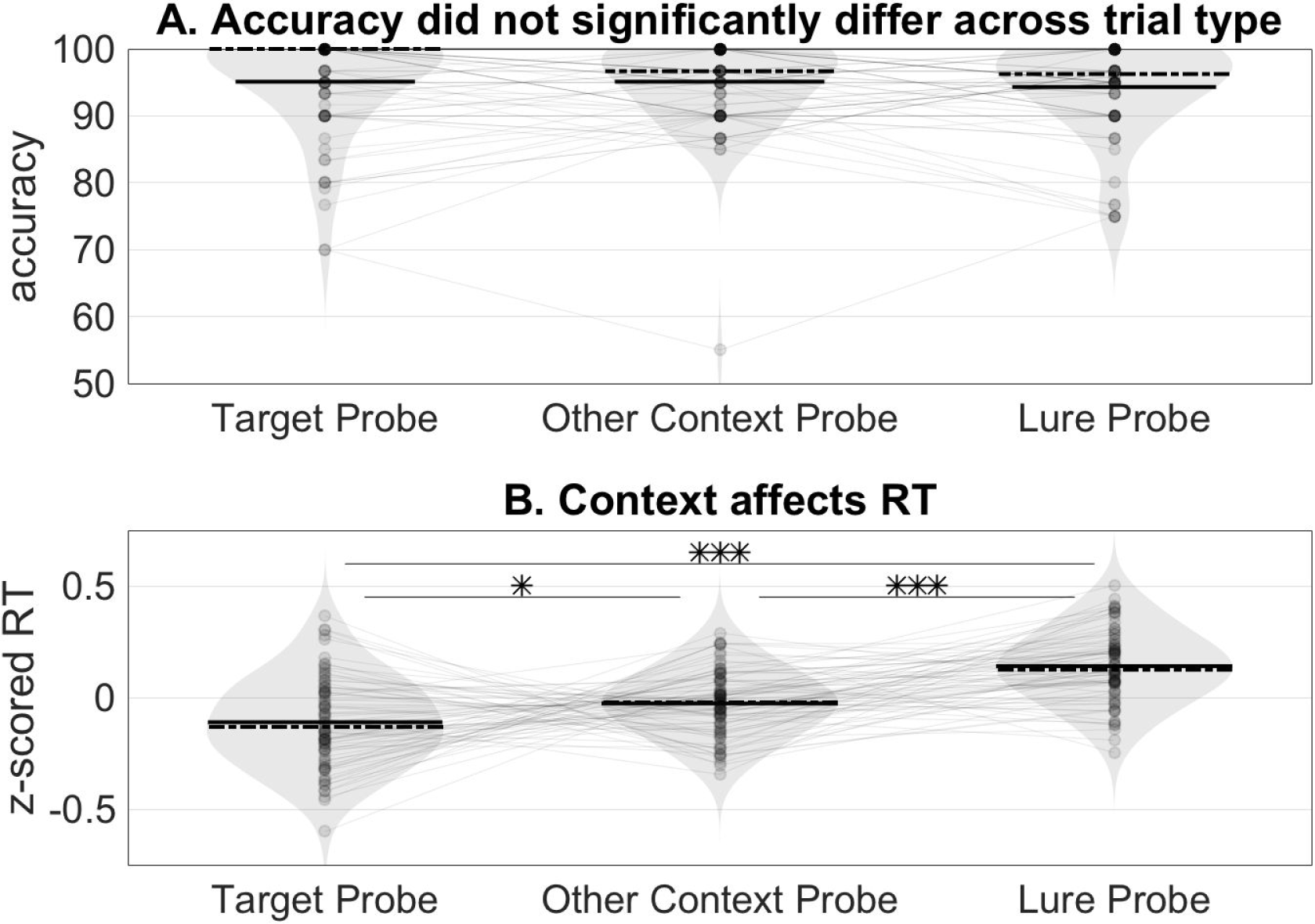
Study 2 results: Response times reflect influence of study context. **A.** For participants with above chance performance (*n* = 80), accuracy was high across all three conditions (mean = 94.84%, SEM = .78%) with no difference in accuracy between *target* (mean = 95.01%, std = 7.37%), *other context* (mean = 95.10%, std = 6.59%), or *lure trials* (mean = 94.31%, std = 6.94%), (*p* > .2 by paired, two-sided t-tests for all pairwise comparisons). Solid lines reflect mean accuracy. Dashed lines reflect median accuracy. **B.** RTs were log transformed and z-scored within-subject to control for individual differences in mean reaction times and non-normal RT distributions. Task irrelevant context information slowed RTs; using paired, two-sided t-tests, we found that participants responded slower to *lure probes* (mean zRT = .14, std = .16) than to *target probes* (mean zRT = −0.11, std = .20; *t*(79) = −6.7603, *p* < .001) or *other context* probes (mean zRT = −0.03, std = .14; *t*(79) = −6.8583, *p* < .001). * = *p* < .05, *** = *p* < .001.

#### Reaction times

We predicted that participants would on average respond fastest to *target probes*, as the probe word would most reliably match the contents of working memory (Figure 3D). In contrast, *non-target probe* trials, in which the probe word did not match any of the targets, would be slower because they required an exhaustive search of the contents of working memory to decide on rejection (a prediction that follows from both serial and parallel models of working memory search — Sternberg, 1969; Ratcliff, 1978).

On *non-target probe* trials, which included *lure* and *other context* probes, participants had to make the same response: to reject the probe word as one of the targets. Thus, any difference in RT between these two trial types could not be attributed to differences in the required response.

Within *non-target probe* trials, we predicted that participants would be slower to respond to *lure* than *other context* probes: If context reinstatement from episodic memory activates trial-irrelevant words from the same context as the target words, lure words can become activated in working memory. If this occurs, activated lure information will match lure probes, increasing uncertainty and slowing “mismatch” responses to these probes. *Other context* probes would not induce such uncertainty, since they would neither match the targets nor would they match reinstated lure information.

RTs were log transformed and z-scored within-subject to control for individual differences in mean reaction times or non-normal RT distributions, however the results reported below are also present in the raw RTs (Supplemental Figure 1).

Using paired, two-sided t-tests, we found that participants responded fastest to *target* probes (mean zRT = −0.11, SEM = .02) compared to *lure probes* (mean zRT = .14, SEM = .02; *t*(79) = −6.7603, *p* < .001; Figure 4B), or *other context* probes (mean zRT = −0.03, SEM = .02; *t*(79) = −2.4133, *p* = .018). Critically, we found participants responded slower to *lure* probes than *other context* probes (*t*(79) = −6.8583, *p* < .001; Figure 4B). The latter is noteworthy as the the only difference between *lure* and *other context* probes is whether the probe word was learned in the same context as the target during the task-irrelevant part of the experiment.

### Experiment 2 discussion

In Experiment 2, participants performed a DNMS task using study words that had previously been associated with one of four separate contexts. The lack of distraction and the relatively short (18 second) delay period were chosen to make it easy for participants to rely solely on working memory to perform the task. Indeed, as has been repeatedly observed in tasks with this kind of structure, accuracy was near ceiling, and did not differ across trial types. However, we observed an effect of encoding context on response times. Specifically, while responses to target probes were faster than responses to both kinds of non-target probes, responses to lure probes — those sharing an encoding context with the target — were slower than responses to probes from any of the other three contexts.

This result is particularly striking because it is in the opposite direction of what would be expected if responses were simply biased towards the more prevalent response type (mismatch). If this were the case, then participants should be faster to respond to lure or other-context probes (⅔ of trials), rather than target probes (⅓ of trials). Instead, the results support the idea that responses may reflect deliberative accumulation of information from working memory, and that this process can be slowed by the intrusion of countervailing information: the context-driven reinstatement of lure words from episodic memory. These reinstatements need not catastrophically interfere with maintenance — rather than occupying discrete “slots” in working memory, they may simply reduce the fidelity of the representation of the target set (e.g., Ma et al., 2014), slowing the integration process without producing an incorrect response.

Note that the same logic should apply irrespective of whether the probe is a lure or an other-context probe — if the correct response is “mismatch”, but (during the delay) participants mentally reinstate the context matching the probe, this should lead to slower RTs to that probe. However, reinstatements of the target-word context should be much more frequent than reinstatements of other contexts, which would explain why responses to lure probes (from the target context) are slower, on average, than are responses to other-context probes.

## Experiment 3

Experiment 2 demonstrated that encoding context has an effect on responses following a delay, even in the absence of distraction. We interpret this result as following from putative episodic memory reinstatements during the delay period. We reasoned that this effect, observed in Experiment 2 as an average across trials, should be determined on a trial-by-trial basis by whether episodic memory reinstatement of the probe context occurred on that trial, as well as which memories were reinstated. To directly test this, in Experiment 3, we had participants perform the same distraction-free DNMS task from Experiment 2 while being scanned using functional magnetic resonance imaging (fMRI), which allowed us to use multivariate pattern analysis (MVPA) to measure the content of memory reinstatement on each trial.

### Methods and materials

#### Participants

40 healthy participants (26 females; ages 18 to 30) were recruited. All participants had normal or corrected-to-normal vision and provided informed consent. The Princeton University IRB approved the study protocol. Exclusion criteria for recruitment included the presence of metal in the body, claustrophobia, neurological diseases or disorders, tattoos above the waist, pregnancy, not speaking English as a native language, and left-handedness. 4 participants were excluded from the final analyses for the following reasons: excessive movement in the scanner — defined as maximal instantaneous displacement larger than 3 mm across any individual scanner run (2 participants), or numerically below-chance accuracy on the DNMS task (2 participants). Data are reported for the remaining 36 participants.

#### Stimuli

The *Fixation* phase used scene and scrambled scene pictures that were not used in any other phase of the experiment. In the *Learning* phase, participants learned four word sets each with its own context picture. The pictures were either faces or scenes. The face pictures were emotionally neutral and of non-famous individuals, taken from the Psychological Image Collection at Stirling University (PICS; http://pics.stir.ac.uk). The scene pictures depicted two natural, non-famous places. One of the faces and one of the scenes were always displayed on the left side of the screen; the other face and other scene were always displayed on the right side of the screen. Thus, each set was associated with one of the following context stimuli: a face on the left, a face on the right, a scene on the left, or a scene on the right. The *Test* phase followed the same DNMS procedure used in Experiment 2. The *Localizer* phase used a different set of scene pictures, along with scrambled scene pictures, neutral faces, and object pictures. All picture stimuli across all tasks were color photos scaled to the same size (500 ×; 500 pixels), equalized for overall brightness, and were displayed 7 degrees from the right or 7 degrees from the left of fixation.

#### Procedure

Prior to the fMRI session, participants practiced the tasks outside of the MRI scanner. Practice consisted of self-paced reading of written explanations of the fixation, context learning, DNMS, and localizer tasks in addition to a fixed number of practice trials of each task. Participants were encouraged to ask questions in case they needed any instruction clarification. After participants reported that they understood the instructions, they completed another practice trial of the context learning task and DNMS task in the scanner.

After practice in the scanner, participants were given 5 minutes of fixation training during which pictures appeared 7 degrees from the right or left of fixation. The goal of this training was to ensure participants perceived the context pictures as lateralized, rather than turning their gaze directly to the picture. We used an Eyelink 1000 eye tracker (SR Research, Ontario, Canada) to give participants real time feedback; if participants looked away from fixation, the images would disappear and an “X” would appear in the center of the screen until fixation was re-established.

After fixation training, participants completed the context list learning and DNMS tasks described in Experiment 2. Trials in which participants did not respond before the 4 second deadline were excluded from analyses, since there was no response time for these trials.

In the final, *Localizer* phase, participants performed a localizer task that was used to discriminate regions of cortex that preferentially process left-and right-lateralized face and scene pictures. In this task, pictures were presented one at a time, and participants were asked to press a key indicating whether the currently presented picture was the same as the one immediately preceding. Pictures were presented in mini-blocks of 10 presentations each. Eight of the images in each block were trial-unique, and two were repeats. Stimuli in each mini-block were chosen from a large stimulus set of pictures not used in the main experiment, and each belonged to one of four categories-faces, objects, scenes or phase-scrambled scenes. and were presented on either the left or right side of the screen. Thus, there were eight different kinds of mini-block: left-face, right-face, left-object, right-object, left-scene, right-scene, left-scrambled, and right-scrambled. Pictures were each presented for 500 m, and followed by a 1.5 second ITI. Participants completed a total of 24 mini-blocks (three blocks per four picture categories presented on either side of the screen), with each mini-block separated by a 12 second inter-block interval.

Finally, after the scanned portions of the experiment had completed, participants remained in the scanner to complete a memory task. Participants were shown each of the 48 words from context learning, one at a time, above all four context pictures, and asked to report both which context was correct and their confidence about that judgement, between one (low confidence) and four (high confidence).

##### Imaging methods

###### Data acquisition

Functional magnetic resonance images (fMRI) were acquired during Phases 2, 3, and 4: context learning, DNMS test, and localizer. Data were acquired using a 3T Siemens Prisma scanner (Siemens, Erlangen, Germany) with a 64 channel volume head coil, located at the Princeton Neuroscience Institute. Stimuli were presented using a rear-projection system (Psychology Software Tools, Sharpsburg, PA). Vocal responses were recorded using a fiber optic noise cancelling microphone (Optoacoustics, Mazor, Israel), and manual responses were recorded using a fiber-optic button box (Current Designs, Philadelphia, PA). A computer running Matlab (Version 2012b, MathWorks, Natick, MA) controlled stimulus presentation.

Functional brain images were collected using a T2*-weighted gradient-echo echo-planar (EPI) sequence (44 oblique axial slices, 2.5 ×; 2.5 mm inplane, 2.5 mm thickness; echo time 26 ms; TR 1000 ms; flip angle 50°; field of view 192 mm). To register participants to standard space, we collected a high-resolution 3D T1-weighted MPRAGE sequence (1.0 ×; 1.0 ×; 1.0 mm voxels).

###### fMRI data preprocessing

Preprocessing was performed using FSL 5.0.6 (FMRIB’s Software Library, www.fmrib.ox.ac.uk/fsl). The first 8 volumes of each run were discarded. All images were skull-stripped to improve registration. Images were aligned to correct for participant motion and then aligned to the MPRAGE. The data were then high-pass filtered with a cutoff period of 128 seconds. 5 mm of smoothing was applied to the data.

###### Region of interest definition

Our anatomical regions of interest were fusiform gyrus, parahippocampal gyrus, and lingual gyrus, based on previous reports of visual category-selective patches of cortex — faces (Kanwisher et al., 1997) and scenes (Epstein & Kanwisher, 1998). We created a bilateral mask combining these three regions that was used for all pattern classifier analyses. Masks were made using cortical parcellation in FreeSurfer with the Destrieux cortical atlas.

###### Multivariate pattern analysis

We extracted the time series of BOLD signal in our anatomical regions of interest during the localizer task and labeled each TR according to the category miniblock to which it belonged. These labeled time series were used to train an L2-regularized multinomial logistic regression classifier (Polyn et al., 2005) to predict the four class labels (left face / right face / left scene / right scene). In our classifier, the probabilities that each class is present do not sum to 1 because we do not assume the categories are mutually exclusive (e.g., we do not assume that the presence of left face evidence necessarily indicates right face absence; Lewis-Peacock & Norman, 2014). To establish the sensitivity of our classifier to the four categories of interest, we performed a leave-one-out cross-validation. First, we split the MRI data from the localizer phase into four runs by time. Then, we trained the classifier on three of the runs, and tested its performance on the fourth, repeating this procedure once using each run as the holdout set. The resulting average performance was significantly above chance (chance = 25.00%, mean = 66.99%, std = 18.30%, t(35) = 14.1419, p < .001; one-sample t-test compared to chance).

To examine how context reinstatements during the DNMS task affected RTs, we divided DNMS trials into 3 time periods: the period when the target words were presented (*target presentation*), the delay period during which participants only saw a fixation cross (*delay period*), and the period during which participants saw the probe word and had to respond (*probe presentation*). To account for the hemodynamic lag, we first shifted our TRs by 5 seconds. Our TRs of interest for each event included TRs from 0 to 6 seconds after each event onset (target presentation, delay period start, probe presentation) plus the shift for hemodynamic lag, with a 1 TR offset between each event in order to minimize contamination of signal between the different periods of interest. The trained classifier was then applied to each volume of activity during these three periods of each trial of the DNMS task. The classifier provided a readout of the probability that the BOLD signal during that volume corresponded to a left face, right face, left scene, or right scene image; we will refer to this as *left/right face/scene evidence.*

**Figure 5.**
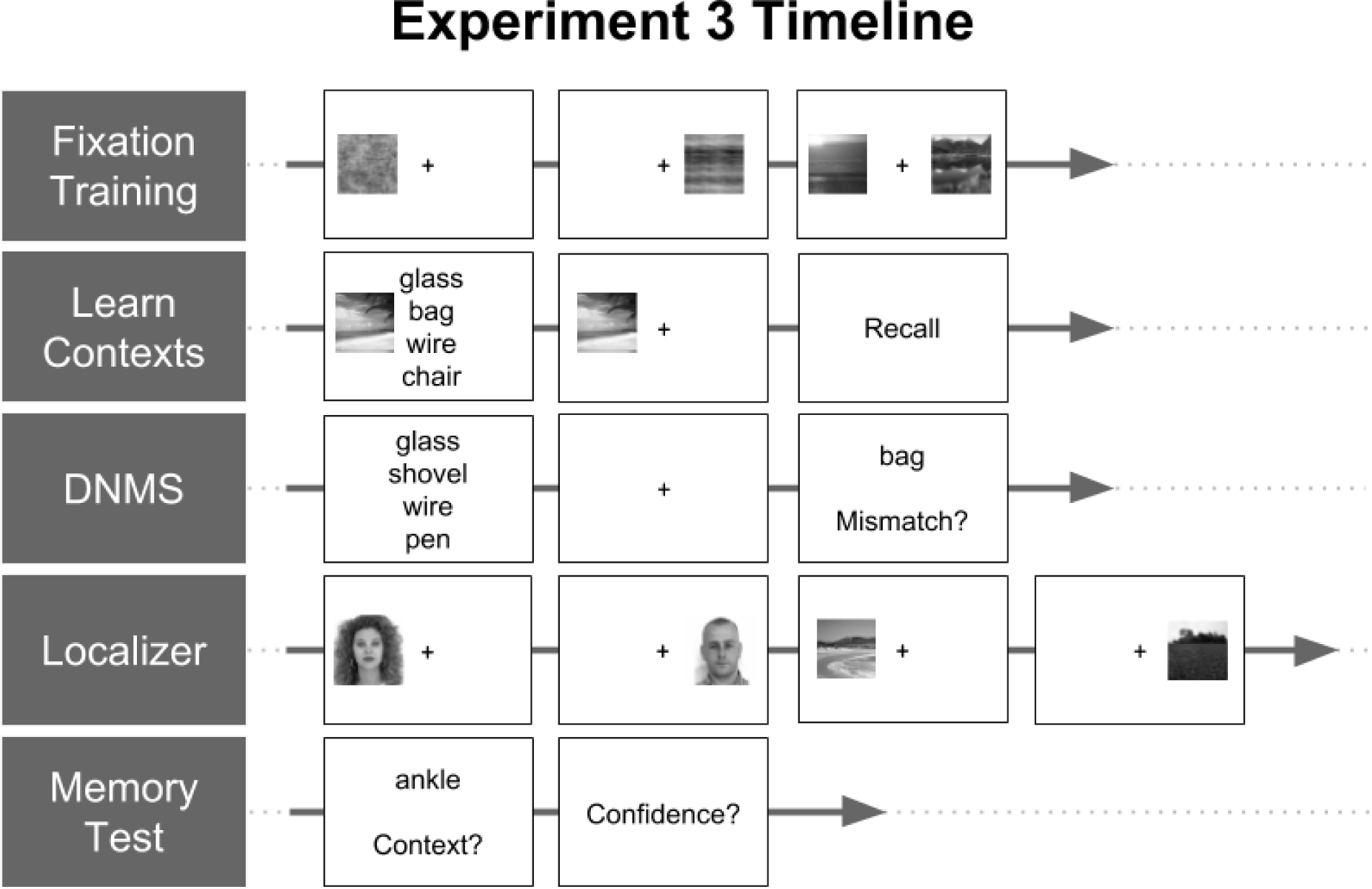
Experiment 3 timeline. We first trained participants to fixate on the center of the screen to ensure that they encoded pictures presented on the left side of the screen as *on the left* and pictures presented on the right side of the screen as *on the right.* Next, participants associated each of four “contexts” (a face presented on the left, a face presented on the right, a scene presented on the left and a scene presented on the right) with a unique set of 12 words. The order in which faces/scenes were displayed on the left/right was randomized across participants. Participants then performed the DNMS task from Experiment 2, after which they performed a one-back localizer task involving blocks of face, scene, object, and scrambled scene images presented on the left/right. Images used during the localizer were distinct from the task stimuli. Finally, participants reported the context with which they thought each word was associated during the initial context-learning phase, in addition to confidence in their report.

### Experiment 3 results

#### Behavioral results

Accuracy for all participants was above the level that would be observed if participants always responded with “mismatch” (66.66%): mean accuracy = 87.27%, SEM = 2.97%. Overall, accuracy on Experiment 3 was significantly lower than mean accuracy on Experiment 2 (unpaired two-sample t-test, t(114) = 3.3797, *p* < .001). As in Experiment 2, accuracy did not differ between the three trial types (Target: mean = 84.44%, SEM = 3.73%; Other-context: mean = 86.25%, SEM = 3.82%; Lure: mean = 87.22%, SEM = 3.76%; paired, two-sided t-tests, all *p* > 0.2).

Due to time restrictions, 3 participants were not able to complete the post-task word/context memory test. The 33 participants who completed the test performed above chance, as a group (chance = 25%, mean accuracy = 41.20%, SEM = 3.33%, t(32) = 4.8648, *p* < .0001, two-sided, one sample compared to chance t-test), and for 25/33 participants individually (proportion *p* <.0001 by binomial test).

As in Experiment 2, we restricted our RT analyses to correct trials only. In contrast to Experiment 2, there was no difference between average RTs in the two mismatch probe conditions (other-context mean log-transformed, z-scored RT = .0311, std = .2313; Lure mean = .0321, std = .1780; *t*(35) = -.0178, *p* = 0.9859; paired sample, two-sided t-test). Consistent with the hypothesis that the slowing on lure trials was driven by reinstated context, RTs *were* different for lure trials on which context-word pairs were later correctly identified in the post-task memory test (β = 11.415, 95% confidence interval = [1.55 21.28], *p* = .02); for each trial type, we estimated the effect of correctly identifying the context belonging to the target words on reaction times using a mixed effects linear regression model for each trial type (Model 1). Remembering the context associated with the target words did not significantly affect reaction times on target or other probe trials, suggesting the slow-down effect of context was selective to trials where context information was misleading (i.e. lure trials).

RT ∼ 1 + TargetMemoryScore * TrialType + (1 | Subject)

Model I: We examine the fixed effects of the different trials (TrialType) and correctly remembering the context belonging to the target word (TargetMemoryScore) on reaction time (RT). We also examine the interaction between the two factors to see whether remembering the target words’ context affects reaction times differently on the different trial types. We control for idiosyncratic individual subject differences by including (1|Subject).

#### fMRI results

We trained an fMRI pattern classifier to discriminate between the four encoding contexts. Then, during the delay period between the presentation of the target words and the probe word, we measured evidence that subjects were reinstating the encoding context associated with the probe words. We predicted that, on mismatch trials (lure and other context probe trials), greater reinstatement of the probe context would cause subjects to be slower to respond, on the assumption that greater activity of the probe word in working memory will make it harder to identify the probe as a mismatch. On target trials, in which the probe word actually was one of the targets, we predicted that reinstating the probe word context would not slow performance.

First, we tested whether context reinstatement lead to slowed responses. We estimated the effect size of probe context reinstatement during our time periods of interest using a mixed effects linear regression model for each trial type (Model 2).

RT ∼ 1 + ProbeContextReinstatementTargetsPresentation + ProbeContextReinstatementsDelay + ProbeContextReinstatementsProbePresentation + (1|Subjcct)

Model 2: We examine the fixed effects of reinstating the probe word’s context during different periods of the DNMS trial on reaction time (RT). ProbeContextReinstatementsTargetsPresentation refers to reinstatements of the probe word’s context during presentation of the targets. The same naming convention applies to probe context reinstatements during the delay period (ProbeContext Reinstatements Delay) as well as during the probe presentation period (ProbeContextReinststatementsProbePresentation). We control for idiosyncratic individual subjeet differences by including (1 | Subjeet).

Supporting our hypothesis, greater evidence for delay-period reinstatement of the probe context was significantly associated with slowed responses on *lure* trials (β = 29.87, 95% confidence interval = [4.39 55.3], *p* = .02), as well as *other-context* probe trials (β = 31.0, 95% confidence interval = [5.1 56.9], *p* = .02; Figure 6A). Further, reinstating the probe context during the delay period on *target* trials did not slow RTs (β = 15.4, 95% confidence interval = [−9.6 40.4], *p* = .23; Figure 6A). (These effects were also unique to delay-period reinstatement; reinstatements during target presentation and probe presentation did not affect reaction times; Supplementary Figure 2).

**Figure 6.**
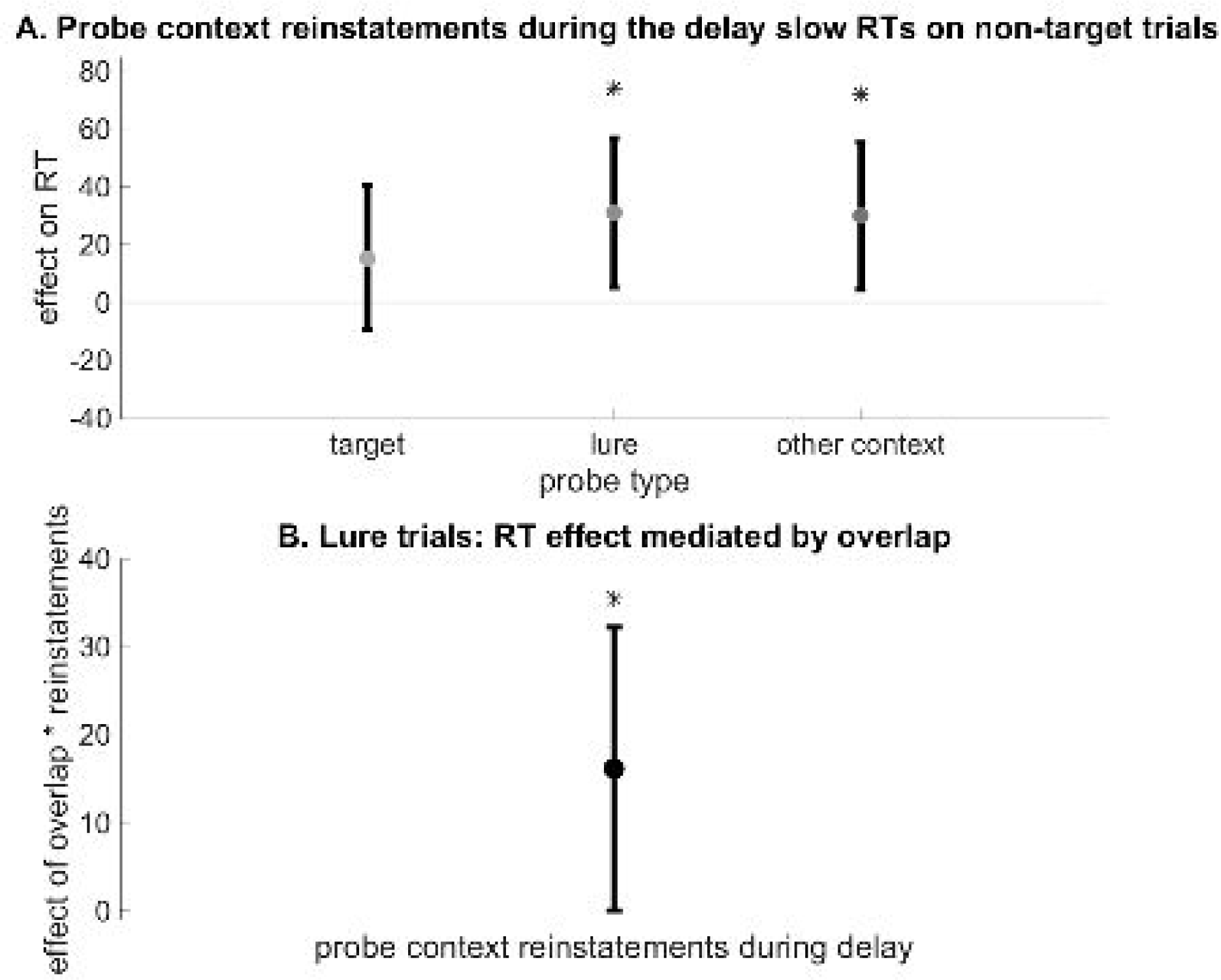
**A.** Greater evidence for delay-period reinstatement of the probe context was associated with slowed responses on lure trials (β = 29.87, 95% confidence interval = [4.39 55.3], *p* = .02), as well as other context probe trials (β = 31.0 95% confidence interval = [5.13 56.87], *p* = .02). Reinstating the probe context during the delay period on target trials did not slow RTs, since these reinstatements did not introduce misleading information into working memory on these trials (β = 15.4, 95% confidence interval = [−9.62 40.42], *p* = .23). *\** indicates *p <* .05. Vertical bars reflect 95% CI. **B.** We predicted that context reinstatements during the delay period would be more likely to slow RTs if the probe word was directly associated not just with the context picture, but also with the target words. For each lure trial, we calculated the number of times the probe word and target words were encountered together during context learning. We found that the more often the probe and targets were encountered together, the more likely participants were to exhibit a slowed response after reinstating the misleading probe context (β = 16.15, 95% confidence interval = [.07 32.22], *p* = .04). This analysis was limited to lure trials because other context probes never overlapped with the targets, and target probes always overlapped with the target words. Vertical bars reflect 95% CI. * indicates *p* < .05.

We hypothesized that this effect would be further mediated by the degree of association between the target and probe words. To test this, we exploited a feature of our experiment that allows us to dissociate between the effects of reinstating pictures versus words: while each word was seen the same number of times with its context picture, there was variation in the number of times each word was presented with another word from the same context during context learning. For each DNMS trial, we computed the number of times the targets and probe were presented together during encoding, a number we called *overlap*; across subjects and trials, overlap scores ranged from 0 to 7 (mean = 3.62, std = 1.50).

We predicted that context reinstatements during the delay period would be more likely to slow RTs if the probe word was directly associated not just with the context picture, but also with the target words (i.e., had higher overlap scores). We used a linear mixed effects regression model to examine how the overlap between the probe and the targets interacted with probe context reinstatements to predict RTs. This analysis was restricted to lure trials only, as, by definition, probes on *other-context* trials were never presented with the target words (Model 3).

RT ∼ 1 + PrthlicContextReinstatementDelay*Overlap + (1 | Subject)

Model 3; We examine the interaction between the number of limes the probe word and target words were presented together (Overlap) and the effect of reinstating the probe word’s context on reaction time (RT). ProbeContextReinstatementsProbePresentation refers to reinstatements of the probe word’s context during the delay. We control for idiosyncratic individual subject differences by including (1 | Subject).

We found a significant interaction between overlap scores and evidence for probe-context reinstatement (β = 16.15, 95% confidence interval = [.07 32.22], *p* = .04; Figure 6B): the more often a given probe overlapped with target words, the more effective reinstatements were at slowing reaction times.

### Experiment 3 discussion

Experiment 3 revealed that memories reinstated during the delay period can alter the contents of working memory, even when these intrusions negatively impact performance on an upcoming match to sample probe.

Using fMRI, we showed that this effect is specific to the degree, timing, and episodic content of the reinstated memories. Namely, disruption results only from context information reinstated during the maintenance period, as opposed to during target or probe presentation. Further, underscoring the episodic nature of these intruding memories, the effect was greater when the potentially misleading words had been presented alongside the target words.

Taken together, these results demonstrate that ongoing episodic memory reinstatement intrudes on working memory maintenance.

## General Discussion

By maintaining a high-fidelity record of recent information, working memory allows us to perform tasks that require accurate storage over short periods of time. However, the presence of distraction or the need to focus on a new task can compromise that record and impair performance. Episodic memory complements these characteristics by storing memories over a longer term, at the cost of reduced fidelity and the risk of retrieval failure (Cohen & O’Reilly, 1996; McClelland et al., 1995; O’Reilly & Rudy, 2001).

While the identification and study of these distinct systems has benefited from efforts to isolate them, it seems unlikely that they would operate entirely independently of one another under natural conditions. Regions that exhibit activity associated with the performance of episodic memory tasks have been observed to be active even during rest, suggesting ongoing reinstatement of episodic memories (Wilson & McNaughton, 1994; Carr et al., 2011; Jadhav et al., 2012). These memory reinstatements can lead to the incidental reinstatement of the context in which the memories were experienced (Bornstein & Norman, 2017). These reinstatements have also been observed to involve coordinated activity across the entire brain, including prefrontal areas associated with working memory maintenance (Miller & Cohen, 2001). Thus, in a manner analogous to externally-driven stimuli, internally-driven reinstatements from episodic memory may also impact representations stored in working memory.

Over a series of three experiments, we tested the hypothesis that episodic memory reinstatement influences performance under task conditions traditionally used to assess working memory maintenance, even in the absence of external interference. In Experiment 1 we showed that, when working memory maintenance is disrupted in a delayed recall task, participants intrude other items from the same context as the studied target items.

Experiment 2 revealed that, even when accuracy is near-ceiling, other measures of performance can detect intrusions from episodic memory. On a delayed non-match to sample task (DNMS) with a distraction-free 18 second delay, participants were slowed in their responding to lure probes — words that shared an encoding context with the target set, but which were not actually members of the target set.

Experiment 3 repeated the DNMS task from Experiment 2. Consistent with the possibility that task-irrelevant context information can affect behavior, we found participants slowed down on lure trials when they had correctly encoded the context belonging to the target words. Using fMRI in Experiment 3 allowed us to investigate the behavioral effects of episodic memory when it was engaged. This analysis revealed that the specific content of episodic memory reinstatement during the delay period predicted the degree of response slowing on that trial.

## The function of replay during working memory maintenance

We have provided evidence that reinstatement of recent experiences from episodic memory has specific, measurable influence on the contents of working memory, even over short delay periods in the absence of explicit interference. Why is working memory influenced by episodic memory reinstatement, even under these conditions? The effect of episodic memory contents on working memory could simply be a side effect, or it could indicate that laboratory tests of working memory maintenance obscure key features of the way that working memory operates in more naturalistic environments. One possibility is that episodic memory is recruited by control mechanisms to “refresh” decaying or disrupted representations.

While some of these reinstatements may be strategically directed recalls in service of maintaining decaying working memory representations, others may instead be ongoing replay of the sort associated with resting-state activity or forward planning (Foster & Wilson, 2006; Tambini et al., 2010; Deuker et al., 2013). On this view, the ability to interact with working memory may be an adaptive feature of resting-state replay from episodic memory — in other words, it may not just sustain, but also transform working memory representations, by integrating information in working memory with information from recent events. That these reinstatements include contextually-related events implies that such an interaction could support rapid, goal-relevant generalizations (Kumaran et al., 2009; Kumaran & McClelland, 2012; Collins & Frank, 2012). The mechanism outlined here both constrains, and expands, that proposal, with potentially broad impacts for the study of memory-guided decision-making.

## Acknowledgements

The authors would like to thank Michael Shvartsman for help with model comparison analysis, as well as Nicholas H. DePinto for technical support with the fMRI scanner and MR-compatible eye tracker. A.N.H. was supported by a National Defense Science and Engineering Grant. A.M.B., A.N.H., K.A.N., and J.D.C. acknowledge support from the Templeton Foundation and the Intel Corporation. The opinions expressed in this publication are those of the authors and do not necessarily reflect the views of the John Templeton Foundation.

## Contributions

A.N.H., A.M.B., and J.D.C. conceived the experiment; A.N.H., A.M.B., J.D.C., and K.A.N. designed the experiments and analyses; A.N.H. wrote the experiment code; A.N.H. ran the experiment; A.N.H. and A.M.B. performed the analyses; A.N.H. and A.M.B. wrote the paper, with input from J.D.C. and K.A.N.

## Data and code availability

The data that support the findings of this study are available on reasonable request from the corresponding author. The data are not yet publicly available because they contain information that could compromise research participant privacy and consent, such as vocal recordings and anatomical brain images. In the near future, they will be anonymized at the level of contemporary best practices and placed in a public repository. All software used to analyze the data are free and publicly available. Standard software packages (SPM8 and FSL 5.0.4) were used for preprocessing the MRI data. The Princeton MVPA toolbox (https://github.com/PrincetonUniversity/princeton-mvpa-toolbox) was used to perform MVPA analyses.

## Supplement

**Supplementary Figure 1.**
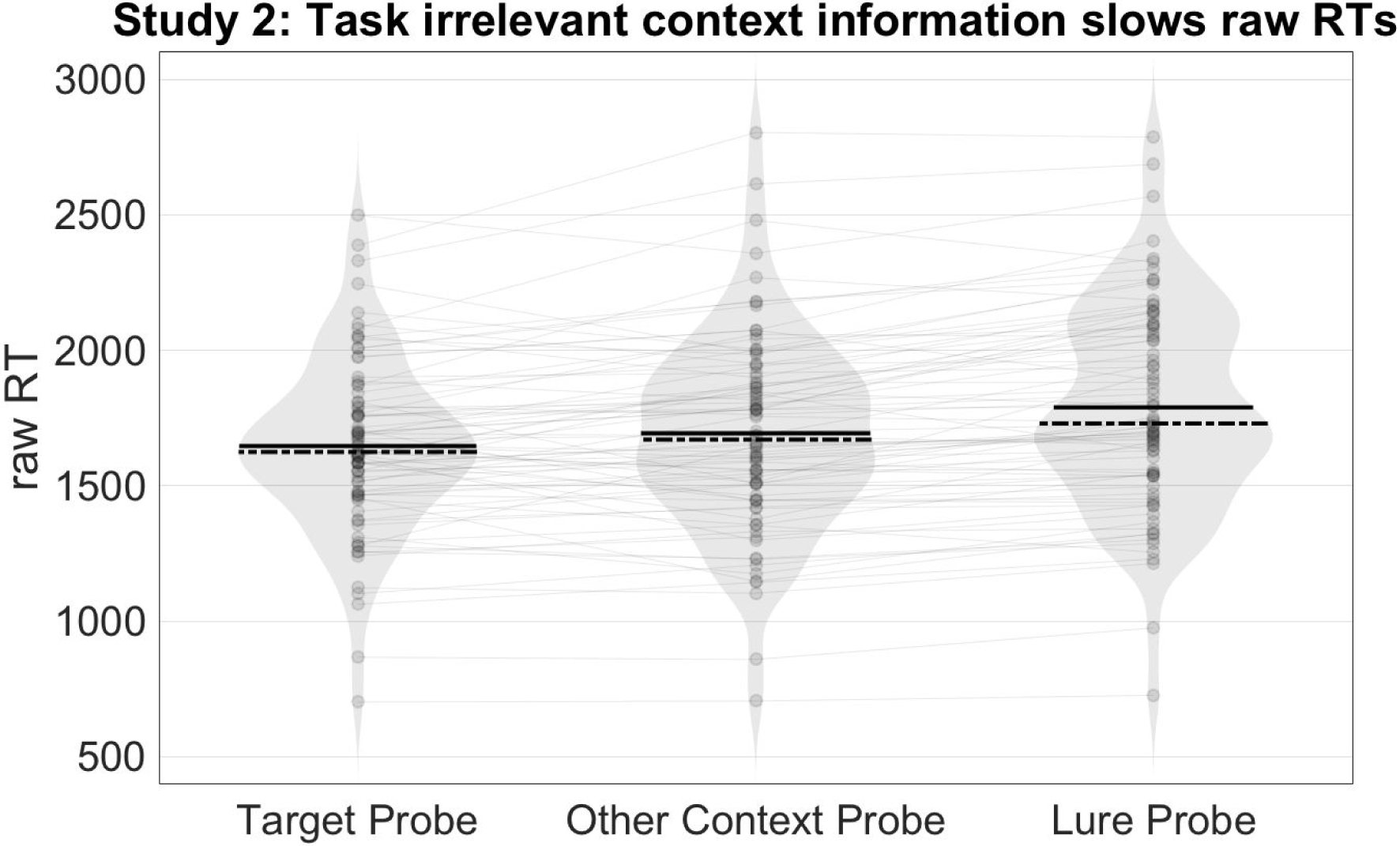
Same analysis as shown in Figure 4B, but using raw RTs. Using paired, two-sided t-tests, we found that participants responded more slowly to *lure probes* (mean RT = 1789.6 ms, std = 381.0 ms), than to *target probes* (mean RT = 1647.2 ms, std = 321.2 ms; *t*(79) = −7.6318, *p* <.001), or *other context* probes (mean RT = 1694.4 ms, std = 366. 9 ms; *t*(79) = −7.0489, *p* < .001). The latter is noteworthy, as the the only difference between these lure and other context probes is whether the probe word was learned in the same context as the target during the task-irrelevant part of the experiment. Solid black lines represent mean RT, dashed lines represent median RT. * = *p* < .05, *** = *p* < .001.

**Supplementary Figure 2.**
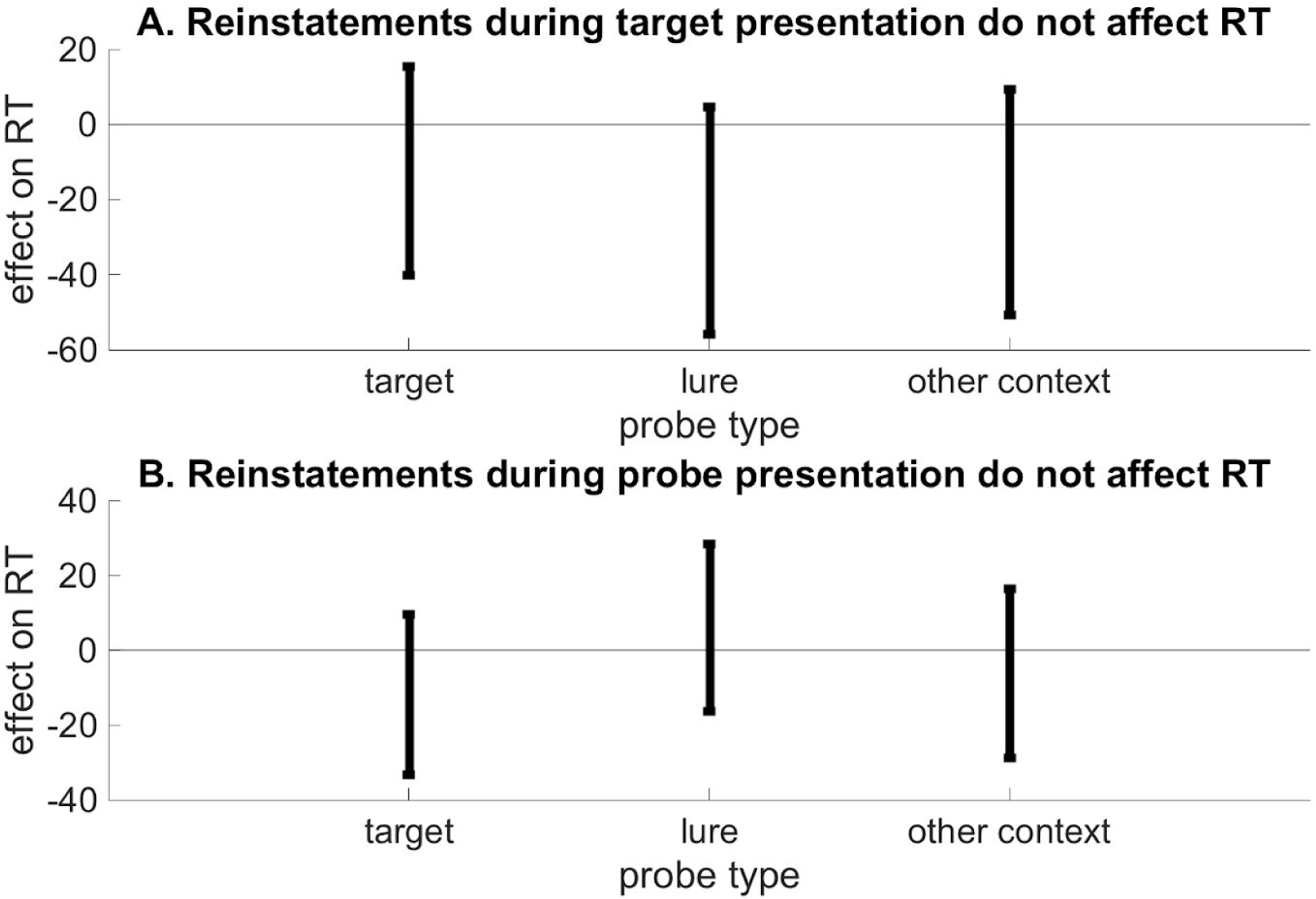
**A.** Using a mixed effects regression model (Model 2, main text), we found there was no effect on reaction times of reinstating the probe word context during the target presentation period. **B.** Results from Model 2 also revealed no effect on reaction times of reinstating the probe word context during the probe presentation period.

## References

Axmacher, N., Mormann, F., Fernández, G., Cohen, M. X., Elger, C. E., and Fell, J. (2007). Sustained neural activity patterns during working memory in the human medial temporal lobe. The Journal of Neuroscience, 27(29), 7807–7816. https://doi.org/10.1523/JNEUROSCI.0962-07.2007

Baddeley, A. (1992). Working Memory, Alan Baddeley. Science, 255(5044), 556–559. https://doi.org/10.1126/science.1736359

Baddeley, A. D., and Hitch, G. (1974). Working memory. The Psychology of Learning and Motivation Advances in Research and Theory. https://doi.org/10.1016/S0079-7421(08)60452-1

Baddeley, A. D., and Hitch, G. J. (2000). Development of working memory: Should the Pascual-Leone and the Baddeley and Hitch models be merged? Journal of Experimental Child Psychology, 77(2), 128–137. https://doi.org/10.1006/jecp.2000.2592

Bornstein, A. M. and Norman, K. A. (2017) Reinstated episodic context guides sampling-based decisions for reward. Nature Neuroscience. https://doi.org/10.1038/nn.4573

Brown, J. (1958). Some tests of the decay theory of immediate memory. Quarterly Journal of Experimental Psychology, 10, 12–21.

Buckner, R. L. (2010). The role of the hippocampus in prediction and imagination. Annual Review of Psychology, 61, 27–48, C1-8. http://doi.org/10.1146/annurev.psych.60.110707.163508

Buckner, R. L., and Carroll, D. C. (2007). Self-projection and the brain. Trends in Cognitive Sciences, 11, 49–57.

Burnham, K.P., Anderson, D.R. (2002). Model selection and multimodel inference: a practical information-theoretic approach, 2nd edn. Springer, New York.

Carr, M.F., Jadhav, S. P., and Frank, L.M. (2011). Hippocampal replay in the awake state: a potential substrate for memory consolidation and retrieval. Nature Neuroscience, 14, 147–153.

Cave, C. B., and Squire, L. R. (1992). Intact verbal and nonverbal short-term memory following damage to the human hippocampus. Hippocampus, 2(2), 151–163. https://doi.org/10.1002/hipo.450020207

Cohen, J.D. and O’Reilly, R.C. (1996). A Preliminary Theory of the Interactions Between Prefrontal Cortex and Hippocampus that Contribute to Planning and Prospective Memory. M. Brandimonte, G.O.

Collins, A. G., Frank, M.J. (2012). How much of reinforcement learning is working memory, not reinforcement learning? A behavioral, computational, and neurogenetic analysis. Eur. J. Neurosci., 35 (2012), pp. 1024–1035.

Einstein and M.A. McDaniel (Eds) Prospective Memory: Theory and Applications, 267-296, Mahwah, New Jersey: Lawrence Erlbaum Associates.

D’Esposito, M., Postle, B. R., and Rypma, B. (2000). Prefrontal cortical contributions to working memory: evidence from event-related fMRI studies. Experimental Brain Research, 133(1), 3–11. https://doi.org/10.1007/s002210000395

Deuker, L., Olligs, J., Fell, J., Krantz, T.A., Mormann, F., Montag, C., Reuter, M., Elger, C.E., and Axmacher, N. Memory consolidation by replay of stimulus-specific neural activity. The Journal of Neuroscience, 33(49), 19373–19383. https://doi.org/10.1523/JNEUROSCI.0414-13.2013

Drachman, D. A., and Arbit, J. (1966). Memory and the hippocampal complex. Archives of Neurology, 15, 52–61. https://doi.org/10.1001/archneur.1964.00460160081008

Epstein, R. and Kanwisher, N. (1998). A cortical representation of the local visual environment. Nature, 392, 598–601.

Foster, D.J., and Wilson, M.A. (2006). Reverse replay of behavioural sequences in hippocampal place cells during the awake state. Nature, 440(7084), 680–683. https://doi.org/10.1038/nature04587

Gershman, S.J., Schapiro, A.C., Hupbach, A. and Norman, K.A. Neural context reinstatement predicts memory misattribution. J. Neurosci. 33, 8590–8595 (2013).

Hannula, D. E., Tranel, D., and Cohen, N. J. (2006). The long and the short of it: Relational memory impairments in amnesia, even at short lags. Journal of Neuroscience, 26(32), 8352–8359. https://doi.org/10.1523/JNEUROSCI.5222-05.2006

Howard, M. W., and Kahana, M. J. (2002). A distributed representation of temporal context. Journal of Mathematical Psychology, 46(3), 269–299. https://doi.org/10.1006/jmps.2001.1388

Hupbach, A., Gomez, R., and Nadel, L. (2009). Episodic memory reconsolidation: Updating or source confusion? Memory (Hove, England), 17(5), 502–510. https://doi.org/10.1080/09658210902882399

Jadhav, S.P., Kemere, C., German, P.W., Frank, L.M.. (2012) Awake hippocampal sharp-wave ripples support spatial memory. Science, 336, 1454–1458.

Kanwisher, N., McDermott, J., Chun, M.M. (1997). The fusiform face area: a module in human extrastriate cortex specialized for face perception. Journal of Neuroscience, 17(11), 4302–4311.

Kumaran, D., Summerfield, J.J., Hassabis, D., Maguire, E. (2009). Tracking the emergence of conceptual knowledge during human decision making. Neuron, 63(6): 889–901.

Kumaran, D., and McClelland, J.L. (2012). Generalization through the recurrent interaction of episodic memories: A model of the hippocampal system. Psychological Review, 119(3), 573–616. https://doi.org/10.1037/a0028681

Lewis-Peacock, J. A., Cohen, J. D., and Norman, K. A. (2016). Neural evidence of the strategic choice between working memory and episodic memory in prospective remembering. Neuropsychologia, 93, 280–288. https://doi.org/10.1016/j.neuropsychologia.2016.11.006

Lewis-Peacock, J.A. and Norman, K.A. (2014). Competition between items in working memory leads to forgetting. Nature Communications, 5(5768).

Logothetis, N. K., Eschenko, O., Murayama, Y., Augath, M., Steudel, T., Evrard, H. C., Besserve, M., Oeltermann, A. (2012). Hippocampal-cortical interaction during periods of subcortical silence. Nature, 491(7425), 547–53. http://doi.org/10.1038/nature11618

Ma, W.J., Husain, M., Bays, P.M. (2014). Changing concepts of working memory. Nature Neuroscience 17(3), 347–356.

Miller, E. K., and Cohen, J. D. (2001). An integrative theory of prefrontal function. Annual Review of Neuroscience, 24, 167–202.

Peterson, L.R., and Peterson, M. J. (1959). Short-term retention of individual verbal items. Journal of Experimental Psychology, 58, 193–198.

Polyn, S. M., Natu, V. S., Cohen, J. D., and Norman, K. A. (2005). Category-specific cortical activity precedes retrieval during memory search. Science (New York, N.Y.), 310(5756), 1963–6. https://doi.org/10.1126/science.1117645

Ranganath, C. (2005). Working memory for visual objects: Complementary roles of inferior temporal, medial temporal, and prefrontal cortex. Neuroscience, 139(1), 277–289. https://doi.org/10.1016/j.neuroscience.2005.06.092

Ranganath, C., and Blumenfeld, R. S. (2005). Doubts about double dissociations between short-and long-term memory. Trends in Cognitive Sciences. https://doi.org/10.1016/j.tics.2005.06.009

Ranganath, C., Cohen, M. X., Dam, C., and D’Esposito, M. (2004). Inferior temporal, prefrontal, and hippocampal contributions to visual working memory maintenance and associative memory retrieval. JNeurosci, 24(16), 3917–3925. https://doi.org/10.1523/JNEUROSCI.5053-03.2004

Ranganath, C., D’Esposito, M., Friederici, A. D., and Ungerleider, L. G. (2005). Directing the mind’s eye: prefrontal, inferior and medial temporal mechanisms for visual working memory This review comes from a themed issue on Cognitive neuroscience Edited. Current Opinion in Neurobiology, 15, 175–182. https://doi.org/10.1016/j.conb.2005.03.017

Repov, G., and Baddeley, A. (2006). The multi-component model of working memory: Explorations in experimental cognitive psychology. Neuroscience, 139(1), 5–21. https://doi.org/10.1016/j.neuroscience.2005.12.061

Rose, N. S., Buchsbaum, B. R., and Craik, F. I. M. (2014). Short-term retention of a single word relies on retrieval from long-term memory when both rehearsal and refreshing are disrupted. Memory and Cognition, 42(5), 689–700. https://doi.org/10.3758/s13421-014-0398-x

Shallice, T., and Warrington, E. K. (1970). Independent functioning of verbal memory stores: a neuropsychological study. The Quarterly Journal of Experimental Psychology, 22(2), 261–273. https://doi.org/10.1080/00335557043000203

Spiegelhalter, D. J., Best, N. G., Carlin, B.P., Van Der Linde, A. (2002). Bayesian measures of model complexity and fit. J. R. Statist. Soc. B, 64(4), 583–639.

Squire, L. R. (1992). Memory and the Hippocampus: A synthesis from findings with rats, monkeys, and humans. Psychological Review, 99(2), 195–231. https://doi.org/10.1037/0033-295X.99.3.582

Sternberg, S. (1969). Memory-scanning: mental processes revealed by reaction time experiments. 57(4), 421–457.

Tambini, A., Ketz, N., and Davachi, L. (2010). Enhanced brain correlations during rest are related to memory for recent experiences. Neuron, 65(2), 280–290. http://doi.org/10.1016/j.neuron.2010.01.001

Tulving, E. (1983). Elements of Episodic Memory. Canadian Psychology, 26(3), 351. https://doi.org/whttp://dx.doi.org/10.1017/S0140525X0004440X

Wickens, D.D., Dalezman, R.E., Eggemeier, F.T. (1976). Multiple encoding of word attributes in memory. Memory and Cognition, 4(3), 307–310.

Wilson, M. (1988). MRC Psycholinguistic Database: Machine-usable dictionary, version 2. 00. Behavior Research Methods, Instruments, and Computers, 20(1), 6–10. https://doi.org/10.3758/BF03202594

Wynn, J.S., Olsen, R. K., Binns, M.A., Buchsbaum, B.R., and Ryan, J.D. (2018). Fixation reinstatement supports visuospatial memory in older adults. Journal of Experimental Psychology: Human Perception and Performance.

Zanto, T. P., Clapp, W. C., Rubens, M. T., Karlsson, J., and Gazzaley, A. (2016). Expectations of task demands dissociate working memory and long-term memory systems. Cerebral Cortex, 26(3), 1176–1186. https://doi.org/10.1093/cercor/bhu307

